# A neural model of modified excitation/inhibition and feedback levels in schizophrenia

**DOI:** 10.1101/2023.04.24.538166

**Authors:** Jiating Zhu, Basilis Zikopoulos, Arash Yazdanbakhsh

## Abstract

The strength of certain visual illusions is weakened in individuals with schizophrenia. Such phenomena have been interpreted as the impaired integration of inhibitory and excitatory neural responses, and impaired top–down feedback mechanisms. To investigate whether and how these factors influence the perceived illusions in individuals with schizophrenia, we propose a two-layer network that can model visual receptive fields (RFs), their inhibitory and excitatory subfields, and the top-down feedback. Our neural model suggests that illusion perception changes in individuals with schizophrenia can be influenced by altered top-down mechanisms and the organization of the on-center off-surround receptive fields. Alteration of the RF inhibitory surround and/or the excitatory center can replicate the difference of illusion precepts between individuals with schizophrenia and normal controls. The results show that the simulated top-down feedback modulation enlarges the difference of the model illusion representations, replicating the difference between the two groups.

---

We propose that the heterogeneity of visual and in general sensory processing in schizophrenia can be largely explained by the degree of top-down feedback reduction, emphasizing the critical role of top-down feedback in illusion perception, and to a lesser extent on the imbalance of excitation/inhibition. Our neural model provides a mechanistic explanation for the modulated visual percepts in schizophrenia with findings that can explain a broad range of visual perceptual observations in previous studies. The two-layer motif of the current model provides a general framework that can be tailored to investigate subcortico-cortical (such as thalamocortical) and cortico-cortical networks, bridging neurobiological changes in schizophrenia and perceptual processing.

## 1 Introduction

Individuals with schizophrenia are less susceptible to illusions, such as the contrast–contrast and apparent motion illusions (Cao et al., 2013;Wurbs et al., 2013;Notredame et al., 2014;King et al., 2017). Lower susceptibility to these illusions could be due to abnormalities in low level integration mechanisms that synthesize inhibitory and excitatory responses of local neurons within primary visual cortex (V1), but also to atypical high-level processes, including reduced top-down influence in perception (Silverstein and Rosen, 2015;Silverstein, 2016;Anderson et al., 2017;Silverstein et al., 2017). However, it is not clear how these factors mechanistically influence the perception of illusions. Development of a neural model, which includes inhibitory and excitatory subfields of receptors, and top-down feedback can provide a platform to characterize functional neural responses based on the visual illusion context, and therefore, can address the challenge of characterizing the impact of each of these three factors on the perception of illusions. Besides schizophrenia, changes of illusion susceptibility have been used to investigate neural processing in other disorders, such as autism spectrum disorder (ASD) (Gori et al., 2016), where neural modeling has been a useful approach to connect underlying neural mechanisms with neurophysiological and perceptual outcomes (Park et al., 2022;Yazdanbakhsh et al., 2023).

It has been shown that people with schizophrenia are less susceptible to the contrast-contrast illusion (Dakin et al., 2005), in which a patch surrounded by a high-contrast background is perceived to have lower contrast than in isolation. It was suggested that this illusion is influenced by contextual surround suppression inhibition (Cavanaugh et al., 2002). However, Barch et al. (Barch et al., 2012) suggested that the illusion percept difference between individuals with schizophrenia and control group may not be significant [reviewed in (King et al., 2017)]; such ambiguity in the findings could stem from the heterogeneity of schizophrenia and its variable impact on visual perception.

Similarly, individuals with schizophrenia have been shown to be less susceptible to apparent motion illusion, in which a pair of separated and alternatively flashing stimuli is perceived as one single moving object (Sanders et al., 2013), likely due to top-down expectations (Edwards et al., 2017). Moreover, neurophysiological evidence supports the role of reduced top–down modulation in decreased susceptibility to hollow-mask illusion in schizophrenia (Dima et al., 2009;Dima et al., 2010;Dima et al., 2011). Based on these findings, it was proposed that the formation of priors (expectations) may affect the top-down process and that the weakened top-down modulation in individuals with schizophrenia can be due to abnormal priors. However, Kaliuzhna et al. (Kaliuzhna et al., 2019) found that there is no evidence for altered formation of priors in schizophrenia.

Given the above broad range of findings, based on variable, often incongruent evidence, a question that needs to be addressed is whether contextual modulation can be a critical factor in reduced illusion susceptibility in schizophrenia. Similarly, what is the role of the top-down feedback in early vision in schizophrenia? We used computational modeling to address these questions by developing a two-layer neural model that can simulate contextual modulation and top-down feedback, in order to probe visual illusion representations in individuals with schizophrenia compared with healthy controls. The two-layer neural model we constructed includes both lateral connectivity and top-down feedback, and therefore supports the investigation of contextual functional interactions. We demonstrate the usefulness of the two-layer neural model in investigating visual processes using two visual illusions, one static (contrast-contrast), and the other dynamic (apparent motion). By considering the model representations of both static and dynamic visual illusions, we can probe the functional interactions between bottom up, top down, and lateral network connectivities.

## 2 Materials and Methods

### 2.1 Model circuit

We developed a two-layer neural model that provides a motif to investigate functional interactions between two hierarchically interconnected brain regions such as thalamic lateral geniculate nucleus (LGN) and primary visual cortex (V1), higher and lower visual cortices, and higher order association and primary cortical areas (Figure1A). In this study, we primarily focus on modeling the visual processing of LGN and V1. However, the emergent properties of the two-layer motif are also applicable to other pairs of interconnected lower and higher visual areas. We can also consider the motif as a unit for feedforward-feedback organization in cortico-cortical or cortico-subcortical communications.

The two-layer motif of our neural model is illustrated in Figure 1B. The output of the neurons in layer 2 is the result of 5 stages (Figure 1C). In stage 1, layer 1 receives the stimulus input and generates its output, which in turn forwards to layer 2 in stage 2. In stage 3, layer 2 feeds back to layer 1. Stage 4 is similar to the feedforward process as stage 1 but takes both the stimulus and the feedback information from stage 3 as the input. Stage 5 is similar to stage 2 and the whole process continues within a feedforward-feedback loop.

**Figure 1.**
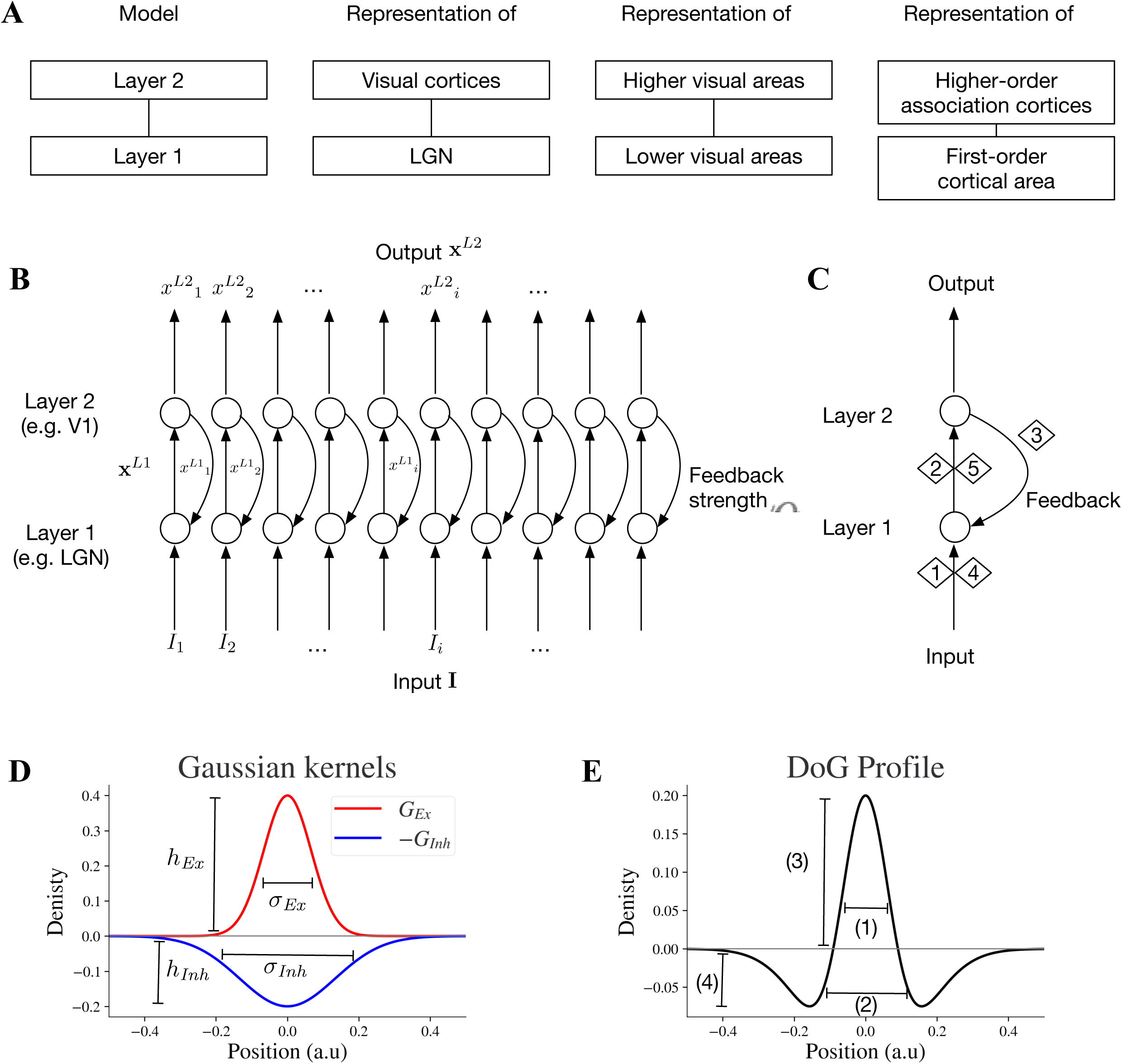
Key specifications and circuitry of the two-layer model. (A) The two-layer model can be considered a general motif applicable to pairs of interconnected subcortical or lower-cortical and higher-order cortical areas. (B) Two-layer network model: the full array of the two-layer network with feedforward and feedback mechanisms. (C) Stages 1,2,4 and 5 are feedforward and stage 3 is feedback. Stage 1 is exposing neurons at layer 1 to input **I** which generates a feedforward output. In stage 2, the output x*^L^*^1^ of layer 1 is the input for layer 2. Stage 4 is the next round of stage 1 with the presence of feedback signal to layer 1, and stage 5 is the next round of stage 2 within the feedforward feedback loop. (D) An excitatory kernel *G_Ex_* and an inhibitory kernel *G_Inh_*. The widths of excitatory and inhibitory subfields depend on their sigma values *σ_Ex_* and *σ_Inh_*, and the amplitudes of excitatory and inhibitory kernels are *h_Ex_* and *h_Inh_*, respectively. (E) Excitatory and inhibitory subfields of model neurons. The baseline positions on the x-axis will be converted to different pixel ranges in simulated visual illusions accordingly. Baseline DoG profile which reflects the spatial spread of the excitatory and inhibitory subfields of the simulated receptive field. The excitatory and inhibitory subfields are modeled by excitatory and inhibitory Gaussian kernels (*G_Ex_*, *G_Inh_*) respectively. 1 reflects the width of excitatory on-center, 2 negative off-surround, 3 peak of on-center, and 4 trough of off-surround.

For each layer, the dynamics of neural activities x over time *t* is formulated within a shunting equation:

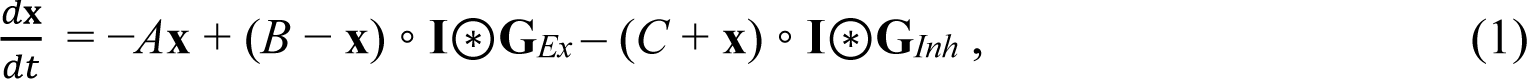

in which **x** is the 1D array (vector) of neural activities, *A* is the decay rate, *B* is the upper limit of **x**, *C* is lower limit of **x**, and **I** is the input array and implemented as a vector (similar to **x**). The symbol ◦ denotes element-wise product and the symbol denotes convolution. Here the convolution takes weighted sum of the input neighborhoods based on the excitatory and inhibitory receptive subfields as inputs to the model neuron array.

**G***_Ex_* and **G***_Inh_* are discretized excitatory and inhibitory Gaussian kernels serving as the excitatory and inhibitory receptive field subfields (i.e., **G***_Ex_* = **G**(*h_Ex_,σ_Ex_*)*/B*, **G***_Inh_* = **G**(*h_Inh_,σ_Inh_*)*/C*, where **G**(*h,σ*) is a vector that discretizes a Gaussian kernel *g*(*k*;*h,σ*)). A Gaussian kernel is formulated as:

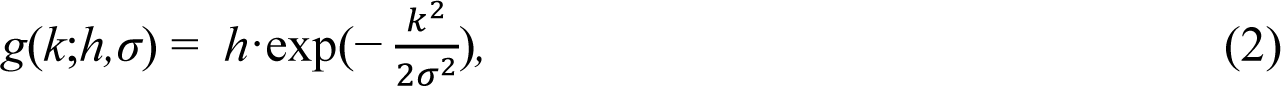

where *k* is the position, *h* represents the amplitude of the peak, and *σ* represents the width of the kernel. The peak amplitude (*h*) and the width of the kernel (*σ*) can be adjusted independently. The excitatory subfield *G_Ex_* is formulated as:

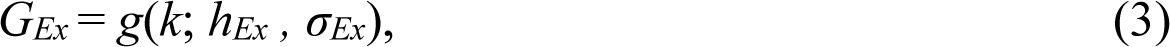

where *h_Ex_* is the peak amplitude of the excitatory kernel and *σ_Ex_* is the excitatory sigma (Fig. 1D).

Similarly, the inhibitory subfield is formulated as

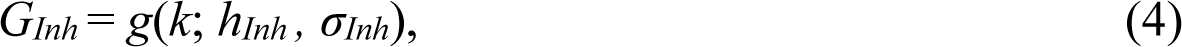

where *h_Inh_* is the peak amplitude of the inhibitory kernel and *σ_Inh_* is the inhibitory sigma (Fig. 1D).

The Gaussian kernel

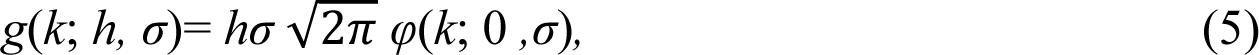

is on the basis of a probability density function *φ*(*k*;0*,σ*) of a normal PDF with mean 0 and standard deviation *σ*:

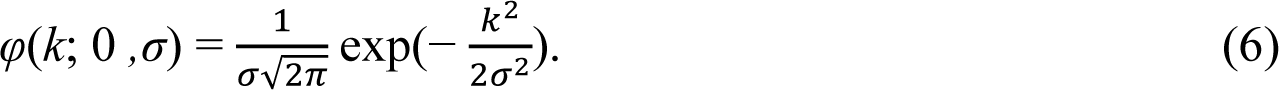

Eq. 5 enables keeping the peak amplitude (*h*) fixed when changing the width of the kernel (*σ*) and vice versa (Zuiderbaan et al., 2012;Anderson et al., 2017).

Modifying the shape of DoG by independent adjustment of *h_Ex_*, *h_Inh_*, *σ_Ex_* and *σ_Inh_* is possible with the above formulation (Eqs. 3 and 4) and implemented throughout the Results section.

In layer 2, neural activities array is **x***^L^*^2^ with input array from **x***^L^*^1^, where **x***^L^*^1^ is the output of layer 1 and **x***^L^*^2^ is the output of layer 2. The feedforward computation for stage 2/5 (Fig. 1C) follows:

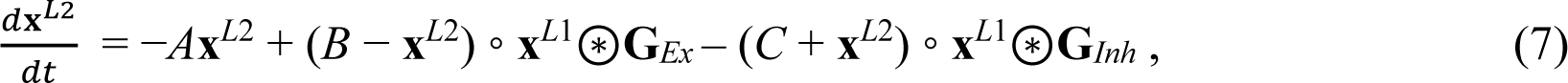

After one round of feedforward-feedback loop, the input to layer 1 is **I**+*α***x***^L^*^2^, where **I** is the stimulus (i.e., illusion) input, and the parameter *α* indicates the connection strength from layer 2 to layer 1 or *feedback strength*. Larger *α* indicates larger feedback, while smaller *α* indicates less feedback. At the start of the first sweep of feedforward signals before feedback signal emergence, **x***^L^*^2^ = 0 at stage 1 without feedback from layer 2. On the other hand, at stage 4 and beyond, when the response from layer 2 is not zero anymore and feeds backs to layer 1, the layer 1 equation is:

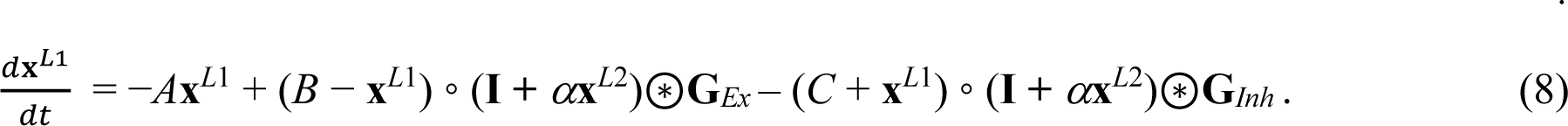

Model parameters are listed in Table 1. Parameter *α* in Eq. 8 determines the feedback strength from model layer 2 to 1. For model excitatory and inhibitory subfields of receptive fields, parameters *h_Ex_* and *h_Inh_* determine peak values of *G_Ex_* and *G_Inh_*, and *σ_Ex_* and *σ_Inh_* determine the widths of *G_Ex_* and *G_Inh_* respectively (Fig. 1D). We investigate how the changes of these parameters affect the model representation of contrast-contrast and AM illusions.

**Table 1:** Abbreviations of symbols used in equations.

| Symbol | Meaning |
| --- | --- |
| $A$ | The decay term |
| $B$ | The upper limit of the model neuron activation |
| $C$ | The lower limit of the model neuron activation |
| $\mathbf{x}$ | Model neurons activation vector with components $x_i$ for each neuron $i$ |
| $\mathbf{I}$ | Input vector with components $I_i$ for each input position $i$ |
| $\mathbf{G}_{Ex}$ | Model representation of excitatory receptive subfield |
| $\mathbf{G}_{Inh}$ | Model representation of inhibitory receptive subfield |
| $G_{Ex}$ | Model excitatory kernel obtained by a function modified from PDF of Gaussian distribution |
| $G_{Inh}$ | Model inhibitory kernel obtained by a function modified from PDF of Gaussian distribution |
| $h_{Ex}$ | Peak amplitude of the excitatory kernel $G_{Ex}$ |
| $h_{Inh}$ | Peak amplitude of the inhibitory kernel $G_{Inh}$ |
| $\sigma_{Ex}$ | Excitatory subfield width (the standard deviation $\sigma$ of the excitatory Gaussian kernel $G_{Ex}$) |
| $\sigma_{Inh}$ | Inhibitory subfield width (the standard deviation $\sigma$ of the inhibitory Gaussian kernel $G_{Inh}$) |
| $\alpha$ | The feedback connection strength from layer 2 (V1) to layer 1 (LGN) (see Equation 3) |

### 2.2 Receptive field

In early stage of visual processing, lateral geniculate nucleus (LGN) neurons respond antagonistically to light in the center and the surround of their receptive fields. Neurons in primary visual cortex (V1) also have oriented on-off center-surround properties (Hubel and Wiesel, 1962). Thus, the receptive field of LGN and primary visual cortex (V1) neurons can be approximated by difference of Gaussian (DoG) kernels in neural models (Layton et al., 2012;Zuiderbaan et al., 2012;Sherbakov and Yazdanbakhsh, 2013;Wurbs et al., 2013;Anderson et al., 2017).

The DoG can be obtained by subtracting a Gaussian with a larger standard deviation from a Gaussian with a smaller standard deviation. The Gaussian with smaller standard deviation, as an excitatory kernel *G_Ex_*, can approximate the on-center of the receptive field (excitatory subfield), whereas the Gaussian with larger standard deviation, as an inhibitory kernel *G_Inh_*, can approximate the off-surround of the receptive field (inhibitory subfield).

The widths of the positive on center and negative off surround and the peak and trough amplitudes of the DoG profile in Fig. 1E depend on the excitatory and inhibitory kernels *G_Ex_* and *G_Inh_* in Fig. 1D.

### 2.3 Simulating contrast-contrast illusion input for the neural model

The input to the neural model is in one dimension and presented as the vector **I**. The model neural response to such one-dimensional input can be characterized by vector **x**. Index *I* indicates the position of each neuron: the model takes the input stimulus with intensity *I_i_* at the position *I* and generates the output response *x_i_* of the neuron *i*.

In order to investigate our model response to the contrast-contrast stimulus (Fig. 2A), we simulated it as an input with a sinusoidal light/dark-gray pattern shown in top of the Fig. 2B, the center of which had lower amplitude than the surround. The simulated 1D input of the contrast-contrast stimulus was a 1D slice from the contrast-contrast stimulus shown in Fig. 2A; its central part had lower contrast than the surround. Although the physical contrast of the inner ringed patch was 40% in Fig. 2A, the surround suppression made it appear less than 40% (Dakin et al., 2005). However, individuals with schizophrenia are less susceptible to the illusion and have more accurate performance than healthy controls for perceiving the actual contrast of the surrounded patch.

**Figure 2.**
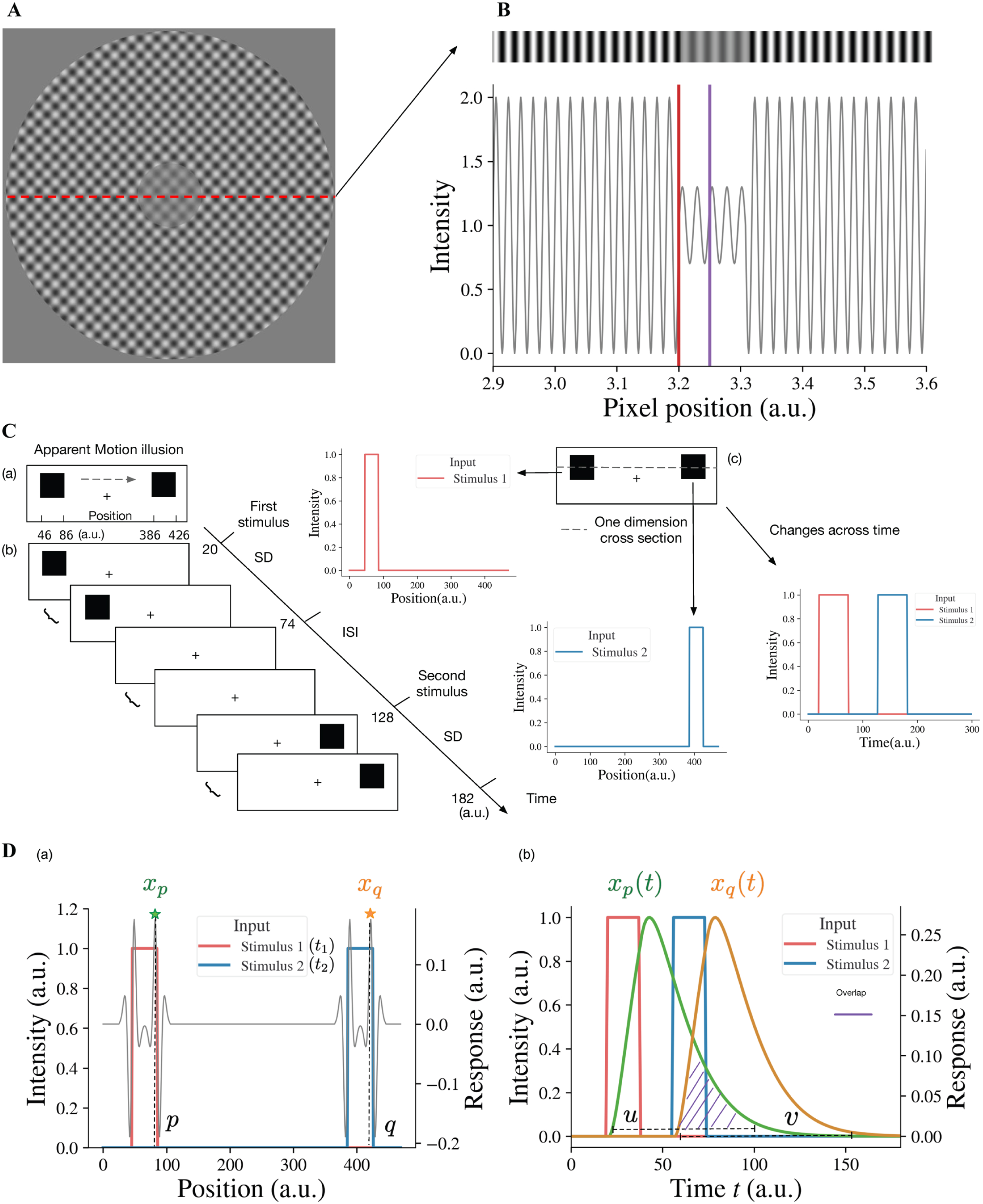
Illusion stimuli as inputs for the model. (A) Sinewave modulation-based contrast-contrast illusion. (B) We take a 1-dimensional (1D) slice from the image in (A) to simulate the input of the model as a 1D gray-scale image shown as a stripe on the top, which shows the visual appearance of the simulated stimulus having lower contrast of light-dark grays at the center, surrounded with higher contrast of light-dark grays. The plot under the gray-scale image is the simulated 1D input of the contrast-contrast stimulus for the neural model and shows the luminance modulation in the form of input intensity modulation across space. The red vertical line is at the left edge of low contrast patch at the middle of the stimulus and the purple vertical line is at the center of the low contrast patch. To avoid the edge effect around the boundaries of the simulated input, padding is applied. Here, the left boundary of this simulated input is located at position 2900 (a.u.). (C) Apparent motion illusion stimulus as an input for the model. (a) Apparent motion (AM) is an illusion of movement created when two adjacent lights flash on and off in succession. (b) Spatial and temporal specification of the apparent motion input to the model. The inter stimulus interval (ISI) has the same duration as the stimulus duration (SD) of the first and the second stimuli. (c) For our model, we consider one dimensional cross-section of the AM stimulus. The first and second stimuli are both spatially and temporally separate, i.e., the first and second offset inputs (lights) flash alternatively (see the panels to the right cued by arrows). (D) The neural model response to the apparent motion stimulus and the model illusion representation quantification by the temporal overlap between the responses to the alternating flashes. (a) The superposition of model response profile to two stimuli, i.e., the spatial profile of the model response to the first stimulus at time *t*_1_, superimposed on the spatial profile of the response to the second stimulus at time *t*_2_ (*t*_2_ *> t*_1_, *t*_2_–*t*_1_ = ISI+SD). *x_p_* denotes the response at position *p*, which is the largest of responses near the right edge of the first stimulus, so does *x_q_* at position *q* of the second stimulus. (b) The changes of responses *x_p_* and *x_q_* over time. *u* is any time point in the time range during which the response *x_p_ >* 0. *v* is any time point in the time range during which the response *x_q_ >* 0. When *u* is bigger than *v*, there is overlap (purple shaded area) between the two response curves. We consider the overlap between the response curve *x_p_*(*t*) and the response curve *x_q_*(*t*), which is the simultaneous occurrence of the two responses, as the continuity representation, i.e., apparent motion, by the model. To quantify the overlap, we consider the response curves *x_p_*(*t*) and *x_q_*(*t*) as two probability density functions after normalizing their surfaces to 1 (see Equation 9).

### 2.4 Model representation of contrast-contrast illusion

To estimate how the surround suppresses the inner contrast, we take the difference between the response to the surround and the response to the center area. A lager value of this difference means a stronger suppression from the surround. Specifically, the illusion representation of the model can be estimated quantitatively as the center-surround response difference, *r* = *s*_1_ −*s*_2_, *s*_1_ being the maximum value of *x_i_^L^*^2^ in the surround region (i.e., to the left of red line, *I <* 3200 (a.u.), in Fig. 2B and *s*_2_ being the maximum value of the simulated responses *x_j_^L^*^2^ to the center area (i.e., around the purple line, 3200 *< j <* 3300 (a.u.), in Fig. 2B. A larger *r* indicates stronger illusion representation by the model and vice versa.

### 2.5 Simulating apparent motion illusion input for the neural model

Apparent motion (AM) is an illusion of movement perception when two or more adjacent lights flash on and off successively appearing as back and forth continuous motion between the two (or more) locations.

Multiple studies have shown that apparent motion is reduced in individuals with schizophrenia (Crawford et al., 2010;Sanders et al., 2013). They tend to report less illusory motion than healthy controls under the same experiment setting.

Consistent with apparent motion stimulus, our simulated input to the model for apparent motion consists of two spatially separated stimuli Fig. 2C(a). After the first stimulus appearance duration (Stimulus Duration (SD), Fig. 2C(b)) the first stimulus disappears, and during interstimulus interval (ISI) there is no visual stimulus. Then the second stimulus appears for the same SD duration. The SDs for the first and second stimuli and ISI are the same. For instance, Fig. 2C(b) shows that the first stimulus appears between 20 time-step (*dt*) and 74 time-step, and the second stimulus appears 54 time-steps after. The presentation duration for both stimuli is also 54 time-steps. The spatiotemporal input to the neural model is a vector **I**(*t*), whose element *I_i_*(*t*) represents the value of input at position *I* at time *t*. The neural model response to the spatiotemporal input **I**(*t*) is a vector **x***^L^*^2^(*t*), where element *x_i_^L^*^2^(*t*) is the response of neuron *I* in layer 2 at time *t*. The model takes the first stimulus *I_ζ_*(*t*_1_) at the position *I* =*ζ* at time instance *t* = *t*_1_ (i.e., first light flash at position *ζ*) and the second stimulus *I_ξ_*(*t*_2_) at the position *I* =*ξ* at time instance *t* =*t*_2_ with *t*_2_−*t*_1_= *b*, and generates the output response *x_i_^L^*^2^(*t*) of each neuron *I* at each time instance *t*; *ξ*–*ζ* is the distance between the two stimuli and *b* is SD + ISI, i.e., the time interval between the onsets of the two stimuli or stimulus onset asynchrony (SOA). For example, *ξ*–*ζ* = 340 and *ζ* is any value between 46 and 86 in Fig. 2C(a), and *b* = 108 and *t*_1_ is any value between 20 and 74 in Fig. 2C(b).

### 2.6 Model representation of apparent motion illusion perception

When the two alternating flashes are presented, the maximum responses are to the edges (Fig. 2D(a)). The peak response to the right edge (at position *p* near 86 in Fig. 2D(a) of the first stimulus is denoted as *x_p_*. The change of the peak response *x_p_* with time is denoted as *x_p_*(*t*) (see the green curve in Fig. 2D(b)). Similarly, the change of the peak response *x_q_* (at position *q* near 426 in Fig. 2D(a)) to the right edge of the second stimulus is denoted as *x_q_*(*t*). The overlap between the two responses is a representation of the continuity of model neural activities between alternating flashes, i.e., the model represents apparent motion (see the purple dashed area in Fig. 2D(b)). The magnitude of overlap can be used as a quantitative measure of model representation of apparent motion.

To calculate the degree of overlap, we consider *x_p_*(*t*) and *x_q_*(*t*) as probability density functions (PDFs) after normalizing their surfaces to 1. Values *u* and *v* in Fig. 2D(b) of the random variables *U* and *V* have *x_p_*(*u*)*/c*_1_ and *x_q_*(*v*)*/c*_2_ as their PDFs, where *c*_1_ and *c*_2_, used for normalization 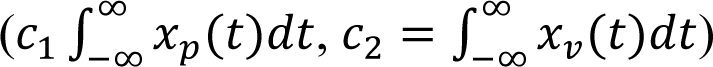 are regarded as modeled representations of the first and second stimuli. The model representation of cross-flashes continuity is the overlap between *x_p_*(*u*) and *x_q_*(*v*), i.e., *u > v* (shaded area in Fig. 2D(b)), which can be determined quantitively by the probability *P*(*U > V*), the probability of simultaneous occurrence of the responses to the first and second stimuli. This model representation, denoted by Φ, for AM illusion perception can be calculated with the convolution of the two probability density functions:

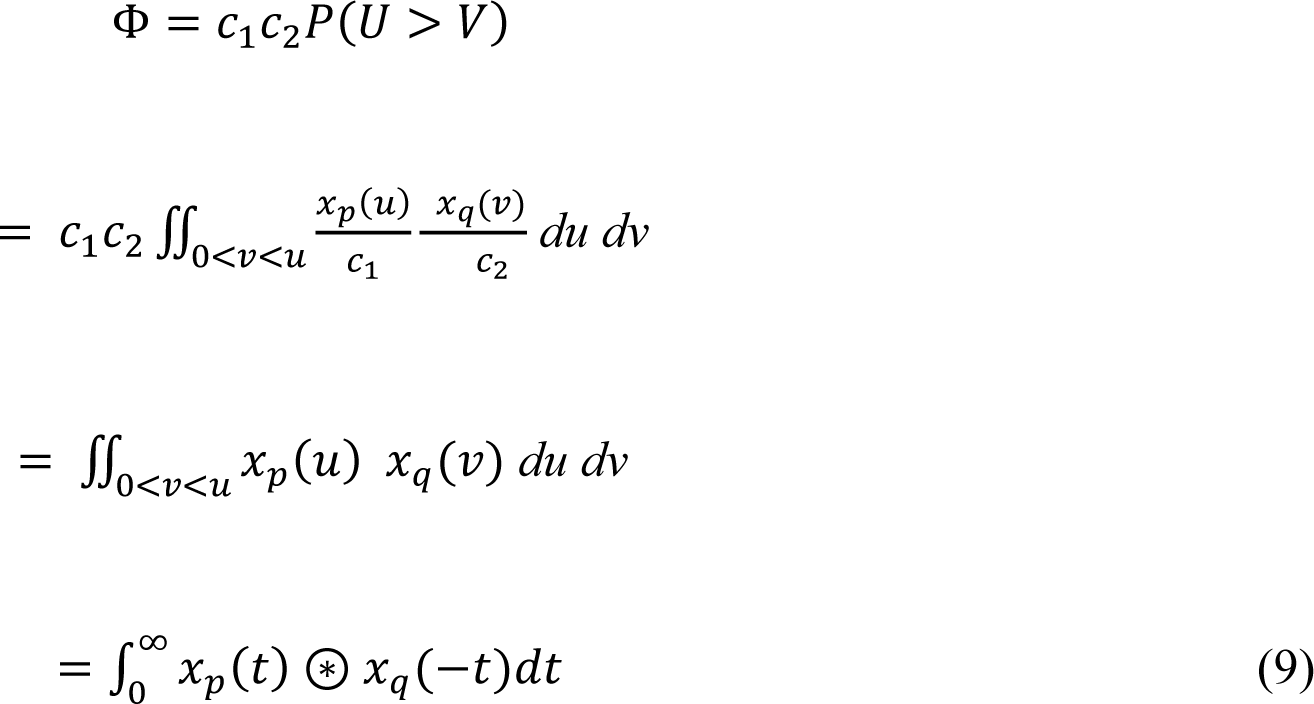

where is again the convolution operation, which is defined as the integral of the product of the two functions after one is reversed and shifted (Bertsekas and Tsitsiklis, 2008). *x_p_*(*u*) is the neuron activity at the time instance *t* = *u* for the first stimulus at the position *i* = *p*, and *x_q_*(*v*) is the neuron activity at the time instance *t* = *v* for the second stimulus at the position *i* = *q*.

## 3 Results

By conducting a thorough parameter search, we investigated the impact of excitation/inhibition width and amplitude ratio changes as well as top-down feedback on the model representation of contrast-contrast and AM illusions. By comparing the existing perceptual data of individuals with schizophrenia with model representations of contrast-contrast and AM illusions, we estimated the range of excitation/inhibition imbalance and the changes of feedback level in the model that can replicate the visual percepts in schizophrenia. Therefore, the key parameter searches we considered were the feedback strength between two model layers and DoG parameters, which were widths and peak amplitudes of excitatory and inhibitory subfields.

When we changed the feedback strength, we kept the DoG profile ratios at the baseline shown in Fig. 3, in which *σ_Ex_/σ_Inh_* =1/2, and *h_Ex_/h_Inh_* = 2. When we changed the DoG parameters, we kept the feedback α at 0.3 and only changed the excitatory and inhibitory subfields in the model V1 (layer 2) by scaling the parameters from the baseline, while the receptive fields in model LGN (layer 1) remained unchanged. In AM simulation, the duration of each stimulus (SDs) was 75, 54, 41, 33, 24, 18, 15, 11, 9, and 6 (in time-steps, each time-step representing 4.44 ms), equivalent to presentation frequencies of 0.75, 1.04, 1.34, 1.7, 2.34, 3.12, 3.75, 4.69, 6.25, or 9.37 Hz, respectively (in a 4×SD cycle). The selected frequencies are consistent with those in Sanders, de Millas, et al. (Sanders et al., 2013). For the parameter search space tested below the model neural representation of contrast-contrast illusion is “*r*”, whereas “Φ” stands for the model neural representation of AM illusion.

**Figure 3.**
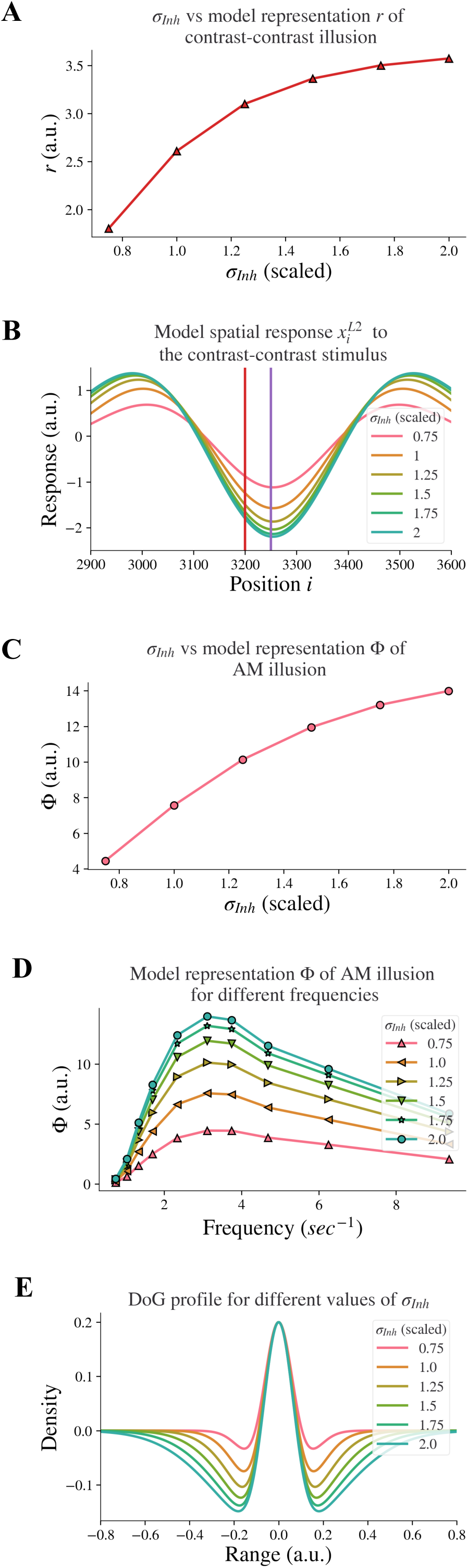
Model response following changes in *σ_Inh_*. The feedback *α* is fixed. (A) The neural representation for contrast-contrast illusion. (B) The model response profile to the contrast-contrast illusion stimulus for different *σ_Inh_*. The red vertical line is at the left edge of the lower contrast patch and the purple vertical line is at the center of the stimulus. (C) The model representation for AM against *σ_Inh_*. (D) The model representation of AM for different frequencies of flashing stimuli, each curve for different *σ_Inh_*. (E) The changes of the DoG profile with the changes of the inhibitory subfield (*σ_Inh_*) used here.

### 3.1 Changes to the inhibitory subfield width led to changes in representation of illusions

Reduced lateral inhibition within V1 in individuals with schizophrenia (Silverstein et al., 2017) is supported by findings of reduced GABA concentration in the visual cortex (Yoon et al., 2010;Yoon et al., 2020). This is in line with reports of reduced inhibition in prefrontal cortices of individuals with schizophrenia (John et al., 2018). Dakin et al. (Dakin et al., 2005) found reduced ‘contrast–contrast’ illusion in individuals with schizophrenia. We investigated whether reduced lateral inhibition caused reduced contrast-contrast illusion. Our results showed that it did: Decreasing the width *σ_Inh_* of the inhibitory subfield in our model led to a reduction of the model representation of contrast-contrast illusion.

We decreased the inhibitory width *σ_Inh_* systematically to model the decreasing of the lateral inhibition. Fig. 3A shows the representation (*r*) of contrast-contrast illusion for a range of *σ_Inh_* values; the scaled value of *σ_Inh_* = 1 is the same value used for the baseline *σ_Inh_* of the DoG profile (see Fig. 1E). Each of the x-axis scaled values in Fig. 3A is the ratio of a *σ_Inh_* value to the baseline value of *σ_Inh_*, showing the key finding that the model representation *r* of contrast-contrast illusion decreased as the inhibitory width *σ_Inh_* decreased. Each value of the model representation *r* in Fig. 3A is the difference between the peak and trough response profile in Fig. 3B. Fig. 3B shows the model response profile *x_i_^L^*^2^ to the contrast-contrast stimulus at each position *i* under systematic change of *σ_Inh_*. The response profiles with the largest, least, and baseline values of *σ_Inh_* are shown in green, pink, and orange, respectively. Each data point in Fig. 3A represents the value of *r* calculated from the corresponding model response profile in Figure 3B, to illustrate the impact of lateral inhibition (*σ_Inh_*) on the model representation (*r*) of contrast-contrast illusion.

Sanders, de Millas, et al. (Sanders et al., 2013) showed that individuals with schizophrenia have impaired AM illusion. Here, we also investigated whether reduced lateral inhibition could lead to reduced AM illusion, and we found that decreasing the inhibitory width *σ_Inh_* in the model also led to reduction of model representation of AM illusion. Fig. 3C shows the model representation (Φ, see Eq. 9) of AM illusion for different values of the inhibitory width *σ_Inh_*: The model representation Φ of AM illusion decreased as the inhibitory width *σ_Inh_* became smaller. Each value of Φ in Fig. 3C is the peak value of its corresponding frequency profile in Fig. 3D. Fig. 3D shows the model representation Φ of AM illusion for different frequencies of the AM stimulus, with each plot for a different value of the inhibitory width *σ_Inh_*. Each plot reached its peak (i.e., the simulated peak representation of AM illusion) around the same frequency (3.12 Hz). This frequency is consistent with the AM frequency in visual experiments yielding peak AM perception (Sanders et al., 2013). With a narrower inhibitory subfield (smaller *σ_Inh_*), the peak drifts to a lower value of the model representation of AM illusion. All the peak values in Fig. 3D are plotted in Figure 3C, which shows the peak representation of AM illusion against the inhibitory width *σ_Inh_*.

Moreover, decreasing the inhibitory width *σ_Inh_* in our model led to a decrease of the inhibitory subfield (or lateral inhibition). Fig. 3E shows the change of the DoG profile under systematic change of *σ_Inh_*. The width of the negative surround shrank as the value of *σ_Inh_* decreased. The pink DoG profile with a smaller *σ_Inh_* showed a narrower inhibition surround than the green DoG profile with a bigger *σ_Inh_*. Furthermore, the trough amplitude of the DoG profile decreased as the inhibitory width *σ_Inh_* decreased. These changes are consistent with (Anderson et al., 2017), where researchers used fMRI and population receptive field (pRF) mapping method to infer properties of visually responsive neurons in individuals with schizophrenia and found that the inhibitory sigma in V1 is significantly smaller in individuals with schizophrenia than in healthy controls. The inter-trough width becomes smaller compared to the DoG profile of V1 in healthy controls (see Figure 4 in (Anderson et al., 2017)). Our results above support the idea that impaired lateral inhibition in V1 would cause reduced contrast-contrast and AM illusions.

**Figure 4.**
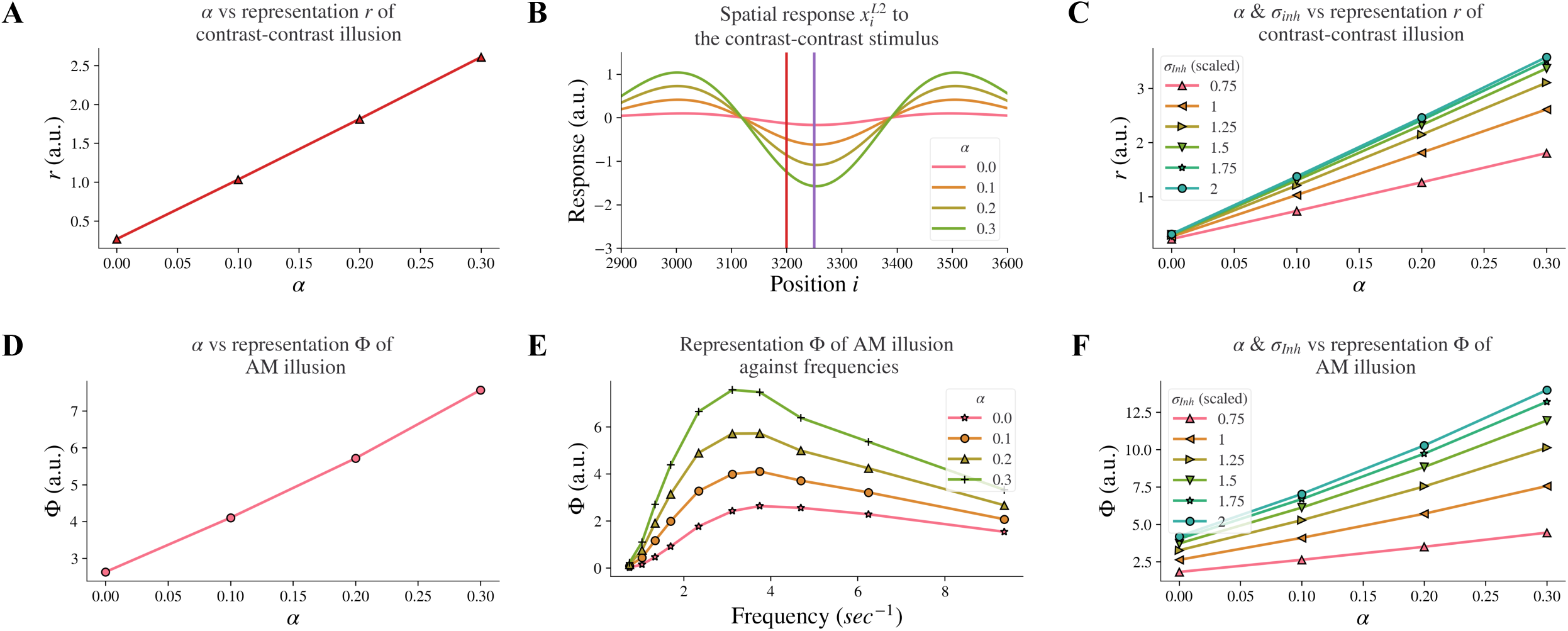
Model responses after changes in feedback strength *α*. (A) The neural representation for contrast-contrast illusion for different *α*. (B) The model response profile to the contrast-contrast illusion stimulus for different *α*. The red vertical line is at the left edge of the lower contrast patch and the purple vertical line is at the center of the stimulus. (C) The model representation for contrast-contrast illusion for different combinations of feedback strength *α* and the width of the inhibitory subfield *σ_Inh_*. (D) The model representation of AM for different feedback *α*. (E) The model representations of AM for different combinations of frequencies of two appearing stimuli and *α*. (F) The model representation of AM for combinations of *α* and *σ_Inh_*.

### 3.2 Weaker feedback strength, led to reduced illusion representation

Individuals with schizophrenia exhibit impaired top-down feedback in visual processing (Gilbert and Sigman, 2007;Silverstein, 2016). In our model, we simulated top-down information flow by controlling the connectivity strength between layers and investigated whether reduced top-down feedback led to reduced contrast-contrast illusion. Our computational model revealed that decreasing the strength *α* of the top-down feedback from layer 2 to layer 1 led to a reduction of model representation of contrast-contrast illusion. Fig. 4A shows the model representation *r* of contrast-contrast illusion for different values of the top-down feedback strength *α*; each data point represents the value *r* from its corresponding plot in Fig. 4B, indicating the key finding that the model representation *r* decreased as the value of *α* decreased. Fig. 4B shows the model response profile *x_i_^L^*^2^ to the contrast-contrast stimulus at each position *i* for different *α* values.

We also investigated whether reduced top-down feedback could influence the contrast-contrast illusion representation as was the case with the reduction of the inhibitory subfield width. Our results showed that it did, but with characteristic differences. The model representation *r* of contrast-contrast illusion was also reduced as the feedback strength *α* and the width *σ_Inh_* of the inhibitory subfield decreased simultaneously (Fig. 4C). It can be observed that the slopes in cool colors (e.g., green) were closer to each other than slopes in warm colors (e.g., pink and orange).

This shows that the increment of the model representation *r* became smaller as *σ_Inh_* increased, while the model representation *r* increased linearly as the feedback *α* increased. As mentioned, reduced top-down feedback led to reduced AM illusion. In Fig. 4D, we can see the model representation Φ of AM illusion for different values of *α*. Fig. 4D presents the peak value of each curve in Fig. 4E, where each curve with a different value of *α* shows the model representation Φ of AM illusion for different frequencies. The model representation Φ of AM illusion went up and down with different frequencies in a way that is consistent with the perceptual data in (Sanders et al., 2013). The peak was lower with a smaller V1 feedback strength *α* and it occurred around the frequency at 3.12 Hz in both our results and the perceptual data. Similar to the results we obtained after decreasing the *σ_Inh_*, reducing feedback strength *α* could also replicate the perceptual results in (Sanders et al., 2013).

When changing the feedback strength *α* and the inhibitory width *σ_Inh_* simultaneously, the model representation of AM illusion had similar trends as that of contrast-contrast illusion. Fig. 4F shows that *σ_Inh_* had a non-linear influence on the model representation Φ while *α* had a linear influence.

All the above results suggest that the disrupted top-down feedback from model layer 2 to layer 1 is also a factor that could underlie the weaker perception of the contrast-contrast and AM illusions in individuals with schizophrenia, but with a different magnitude of impact compared to the disruption in the width of the inhibitory subfield.

### 3.3 Changes in the width of the excitatory subfield had variable impact on illusion representation

Changes in excitation, in particular increases in activity of excitatory neurons, have been associated with schizophrenia. Increases in activity of excitatory neurons in schizophrenia can come about due to decreased excitatory drive on fast-spiking cortical inhibitory interneurons (Chung et al., 2016), or an overactive dopaminergic system (Gracitelli et al., 2015;Howes et al., 2015;Silverstein and Rosen, 2015). However, there is little evidence showing how changes in excitatory drive through decreased inhibition or excess dopamine can influence visual receptive fields. To test the hypothesis that an enlarged excitatory subfield could reflect changes in excitation, we investigated whether increasing the excitatory subfield width would reduce contrast-contrast illusion. We found that this was dependent on whether the excitatory subfield width was large or small.

By increasing the width *σ_Ex_* of the excitatory subfield in the range of larger values (here 0.75-1.5) in the model, we observed a reduction of the model representation *r* of contrast-contrast illusion (see the right panel in Fig. 5A). Figure 5B shows the model response profile *x_i_^L^*^2^ to the contrast-contrast stimulus at each position *i* under systematic change of the *σ_Ex_* in the range of larger values. However, an opposite trend was observed when the excitatory width *σ_Ex_* was in the range of smaller values (0.25-0.75). Increasing the smaller *σ_Ex_* increased the model representation *r* of contrast-contrast illusion (see the left panel in Figure 5A).

**Figure 5.**
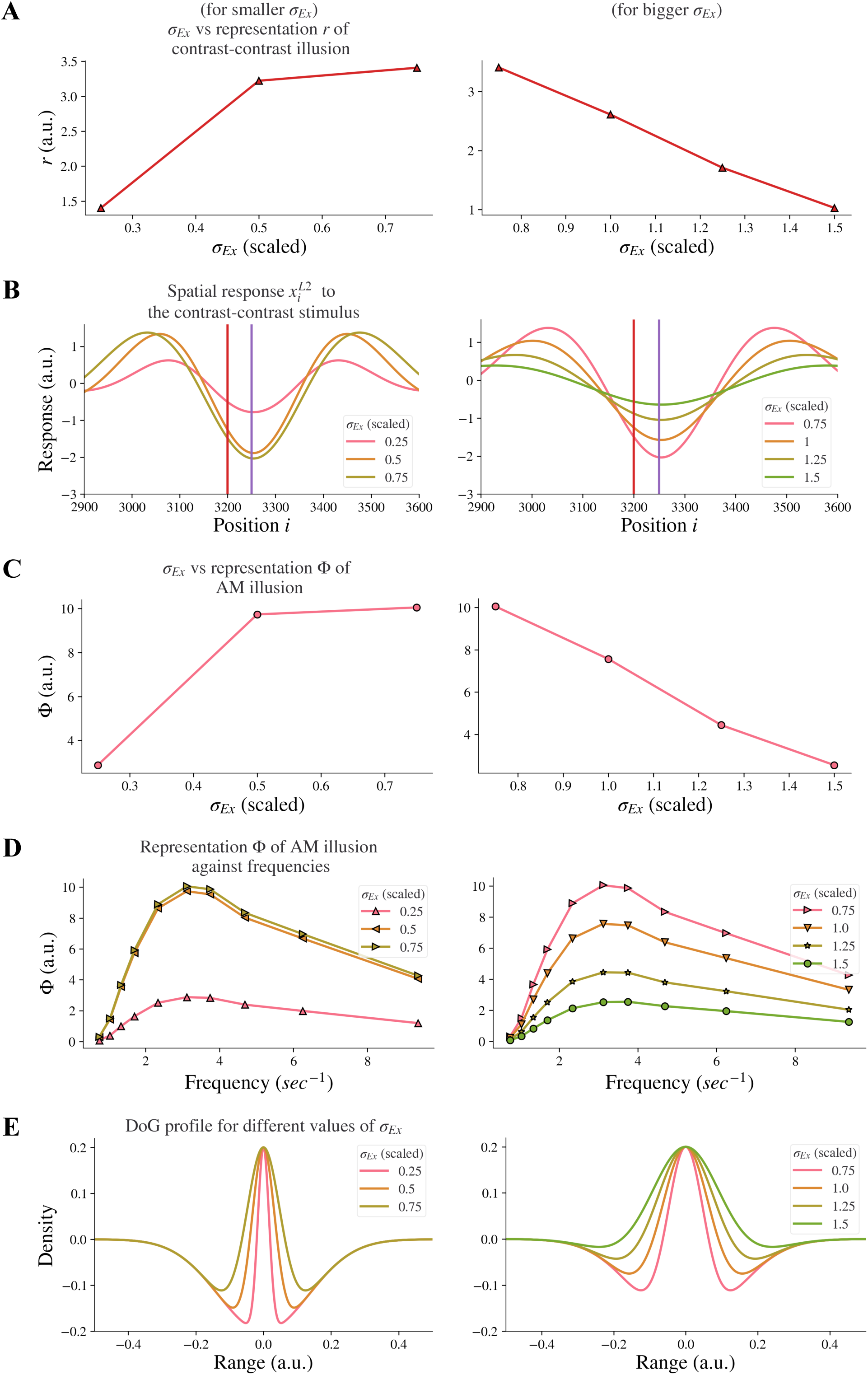
Model representations of illusions for different excitatory width *σ_Ex_*. Results for smaller *σ_Ex_* and bigger *σ_Ex_* are on the left and right panels respectively. (A) The neural representation for contrast-contrast illusion perception. (B) The model response profile for the contrast-contrast illusion. The red vertical is at the left edge of the lower contrast patch and the purple vertical line is at the center of the stimulus. (C) The model representation for AM illusion against each *σ_Ex_*. (D) The model representation for AM illusion against different combinations of *σ_Ex_* and frequencies of the two flashing stimuli. (E) The changes of the DoG profile with the changes of the excitatory subfield *σ_Ex_*.

Similarly, we investigated whether increasing the excitatory subfield width would reduce AM illusion. The results for the AM illusion were consistent with the contrast-contrast illusion: changes varied depending on whether the excitatory subfield width values were within a low or high range.

By increasing the excitatory width *σ_Ex_* in the range of larger values (0.75-1.5) in the model, we observed a reduction of the model representation Φ of AM illusion (see the right panel in Fig. 5C). On the contrary, increasing the excitatory width *σ_Ex_* in the range of lower values (0.25-0.75) increased the model representation Φ of AM illusion (see the left panel in Fig. 5C). Figure 5C presents the peak value of each curve in Fig. 5D. The range of scaled *σ_Ex_* applied here kept the DoG profile in an on-center off-surround shape (Fig. 5E).

Our results suggest that the model base for healthy controls might be at the scaled value of the excitatory width *σ_Ex_* around 0.75, and that individuals with schizophrenia with decreased excitatory drive on fast-spiking interneurons or overactive dopaminergic system could be modeled by scaled value of *σ_Ex_* greater than 0.75. Furthermore, such nonmonotonic changes of illusion representation when the excitatory width *σ_Ex_* increased indicate that the excitatory width *σ_Ex_* may not be the dominant factor that caused the resistance to the illusions in individuals with schizophrenia.

### 3.4 Changes in the inhibitory subfield amplitude did not affect illusion representation

We additionally investigated whether reducing the amplitude *h_Inh_* of the inhibitory subfield, as an alternative approach to model the reduction of the inhibitory subfield, could have the same effect as reducing the width *σ_Inh_* of the inhibitory subfield. Our results showed that reducing the amplitude *h_Inh_* of the inhibitory subfield had an opposite influence on the model representation of illusions compared to the reduction of the width of the inhibitory subfield. We found that reducing the inhibitory amplitude *h_Inh_* did not reduce the model representation *r* of contrast-contrast illusion (see Fig. 6A). Instead, Fig. 6A shows that the model representation *r* of contrast-contrast illusion increased as the value of *h_Inh_* decreased. Each data point in Fig. 6A represents the value of *r* calculated from its corresponding *r* value of each curve in Fig. 6B. Similarly, we found that reducing the inhibitory amplitude *h_Inh_* did not reduce the model representation Φ of AM illusion. Fig. 6C shows that the model representation Φ of AM illusion increased as the value of *h_Inh_* decreased. All the peak values in Fig. 6D are plotted in Fig. 6C. Fig. 6E shows that the on-center area of the DoG profile became larger as the inhibitory amplitude *h_Inh_* decreased.

**Figure 6.**
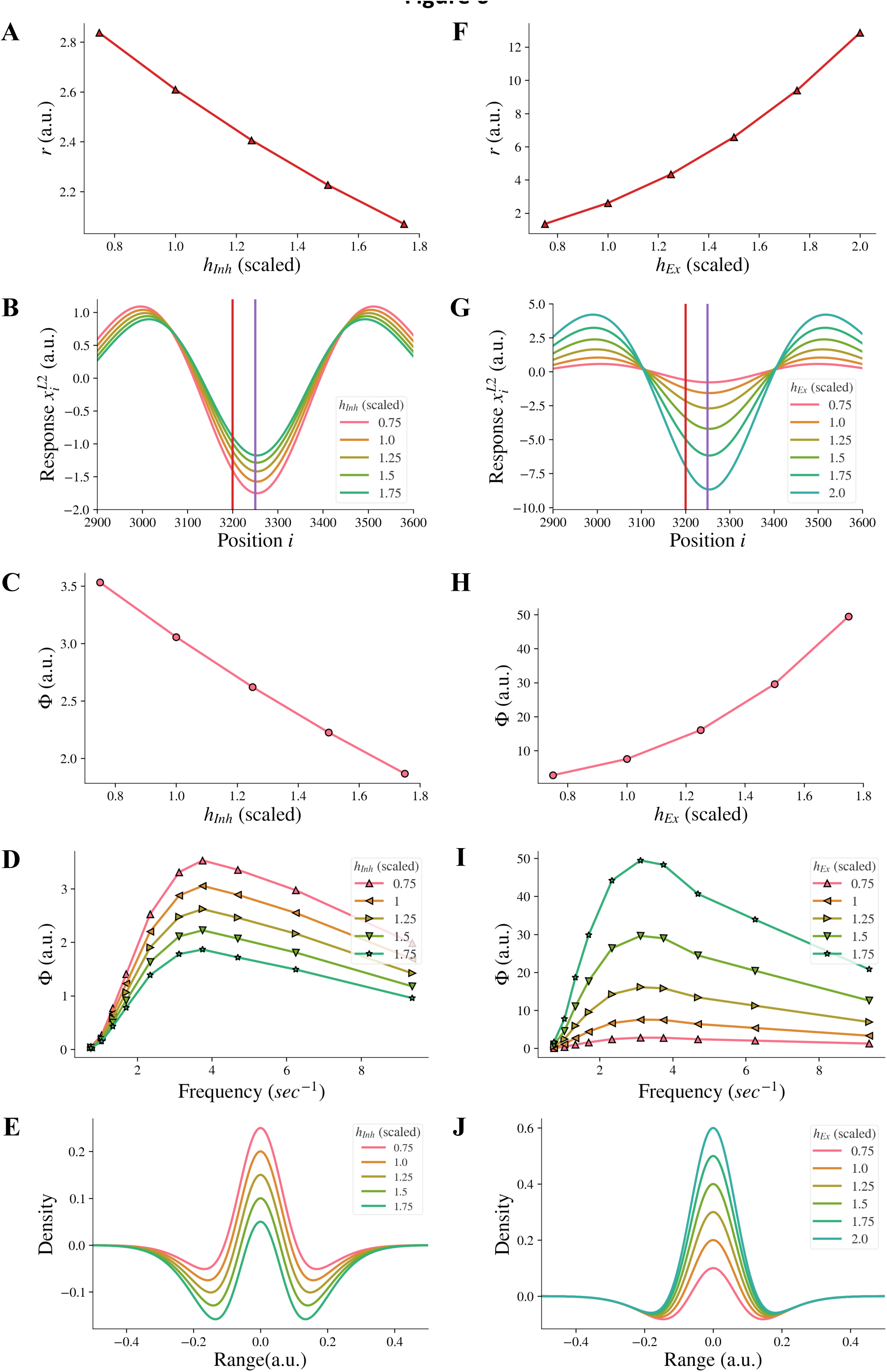
Model representations of illusions for different inhibitory amplitude *h_Inh_* and excitatory amplitude *h_Ex_*. (A) The model representation for contrast-contrast illusion. (B) The model response profile to the contrast-contrast illusion stimulus. The red vertical line is at the left edge of the lower contrast patch and the purple vertical line is at the center of the stimulus. (C) The model representation for AM illusion against each inhibitory amplitude *h_Inh_* . (D) The model representations for AM illusion against different frequencies of two flashing stimuli and *h_Inh_*. (E) The changes of the DoG profile with the changes of the inhibitory amplitude (*h_Inh_*). F-J are equivalent of A-E by replacing changes of *h_Inh_* with changes of *h_Ex_*.

We also examined whether increasing the amplitude *h_Ex_* of the excitatory subfield, as an alternative approach to model the increase of the excitatory subfield, could have the same effect on the illusion representation as increasing the excitatory width *σ_Ex_* in a range of higher values. We found that increasing the excitatory amplitude *h_Ex_* did not reduce the model representation *r* of contrast-contrast illusion. Fig. 6F shows that the model representation *r* of contrast-contrast illusion increased as the value of *h_Ex_* increased. Each data point in Fig. 6F represents the value of *r* calculated from its corresponding *r* value of each curve in Fig. 6G. Similarly, we found that increasing the excitatory amplitude *h_Ex_* did not reduce the model representation Φ of AM illusion. Fig. 6H shows that the model representation Φ of the AM illusion increased as the value of *h_Ex_* increased. All the peak values in Fig. 6I are plotted in Fig. 6H. Fig. 6J shows that the on-center area of the DoG profile became larger as the excitatory amplitude *h_Ex_* increased.

### 3.5 The effect of simultaneously reduced widths of inhibitory and excitatory subfields

When the inhibitory width *σ_Inh_* and the excitatory width *σ_Ex_* changed simultaneously, the contribution of the change of *σ_Ex_* to the model representation was not affected by the change of *σ_Inh_* and vice versa, i.e., there was no interaction, and each factor had its own independent effect on the representation. The model representation *r* of contrast-contrast illusion against the excitatory width *σ_Ex_* is presented in Fig. 7A, with each curve for a different value of the inhibitory width *σ_Inh_*. When the inhibitory width *σ_Inh_* decreased, the curve of the model representation *r* moved down, and the slope remained almost the same. When the excitatory width *σ_Ex_* decreased, the value of the model representation *r* increased monotonically, and the change rate was almost the same for different values of the inhibitory width *σ_Inh_*. We saw the same pattern for the model representation Φ of AM illusion (Fig. 7B).

**Figure 7.**
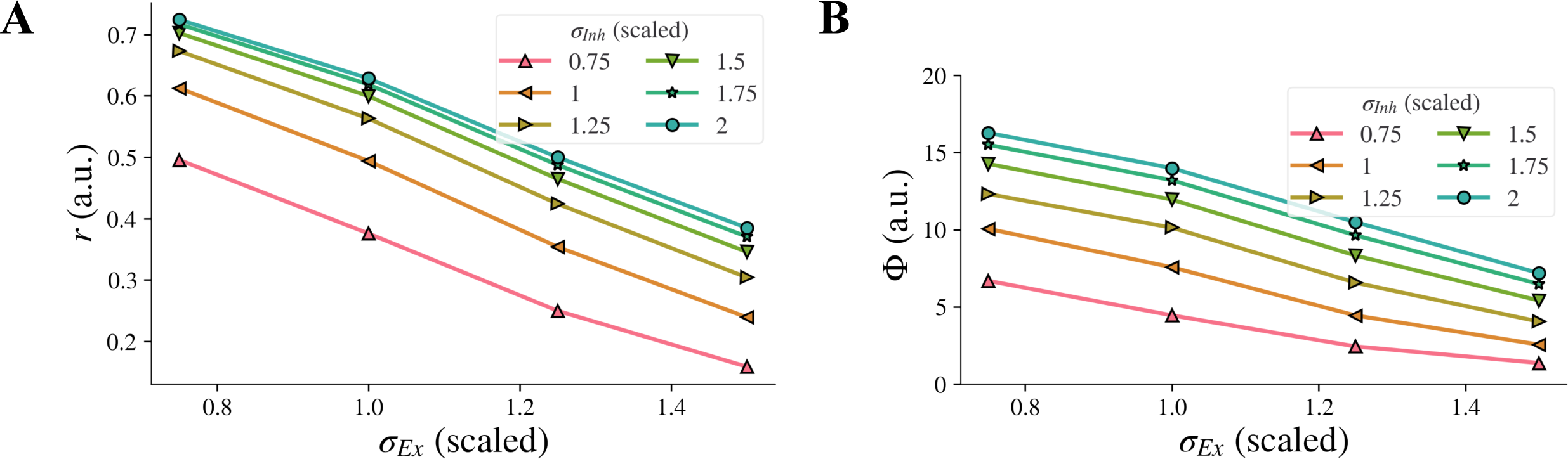
Model illusion representation for combined changes of *σ_Inh_* and *σ_Ex_*. Model representation for contrast-contrast (A) and AM (B) illusions. In both cases, decrease of *σ_Inh_* and simultaneous increase of *σ_Ex_* led to decreased illusion representation.

## 4 Discussion

We constructed a two-layer neural model that can replicate the perceptual results in contrast-contrast and AM illusions, as elaborated below. We observed reduction of simulated illusion representation when changing the model parameters independently or combined, including decrements of the width of the inhibitory surround, decrements of the top-down feedback, and increments or decrements of the width of the excitatory subfield (Fig. 8).

**Figure 8.**
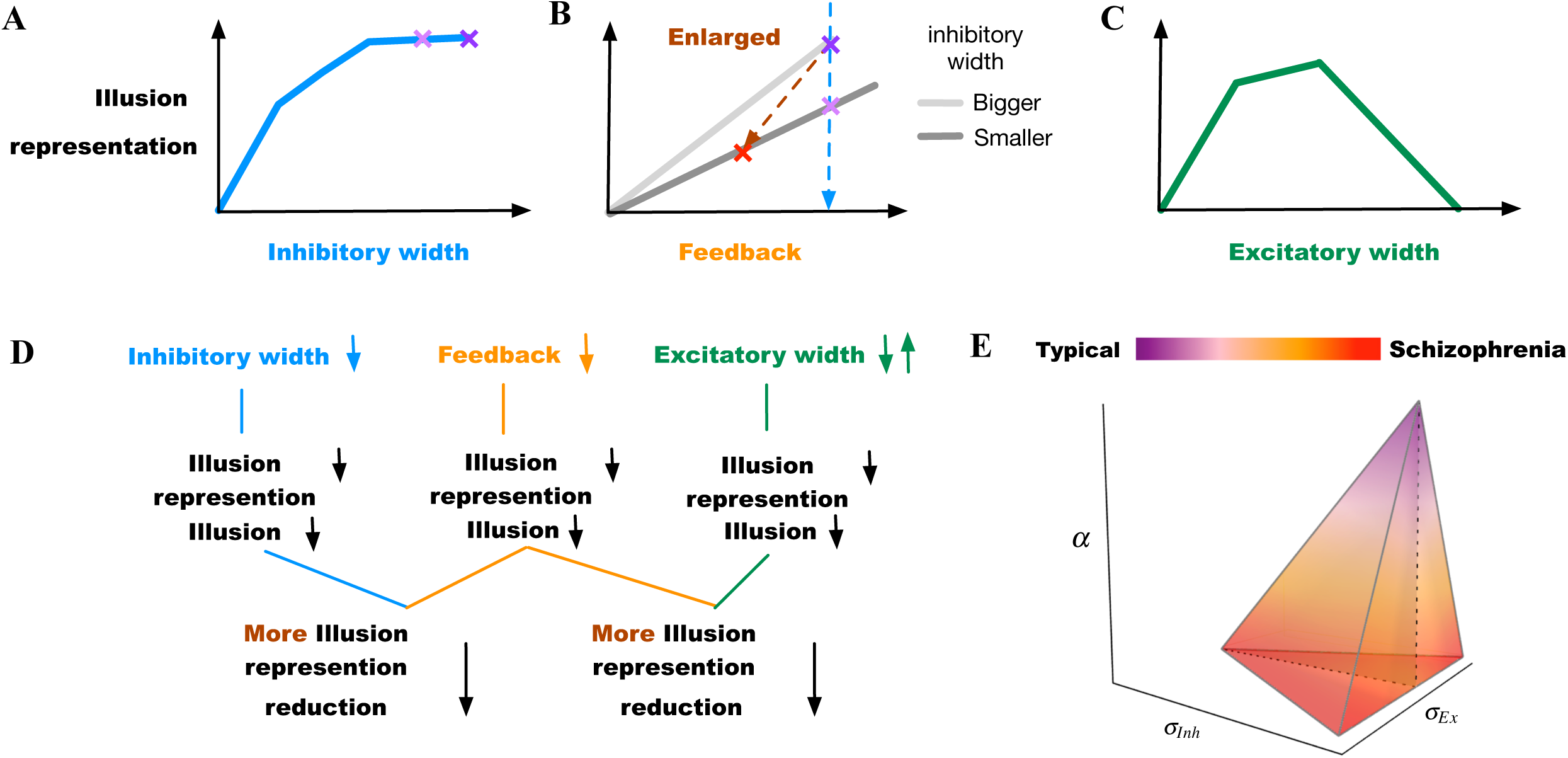
Summary of findings. (A) Model illusion representation reduces as the width of the inhibitory surround decreases. The two purple “x” marks indicate little change within the upper range of inhibitory width values. (B) Model illusion representation reduces as the feedback decreases. The light purple “x” mark indicates the illusion change when the width of the inhibitory surround decreases (the same as in (A)). The red “x” mark indicates a greater reduction of illusion representation, when both the inhibitory width and the feedback decrease simultaneously. (C) Model illusion representation reduces as the width of the excitatory subfield increases or decreases depending on different starting points. (D) Summarizing independent possibilities for reduced model illusion representation and their combination resulting in further reduced representation. (E) Transition from typically developed individuals to those with schizophrenia within *α*, *σ_Inh_*, and *σ_Ex_* (feedback, inhibitory, and excitatory subfields) space. The schizophrenia region is within ranges of reduced feedback and inhibition with either reduced or increased excitation highlighting neurobiological heterogeneity.

### 4.1 Comparison with behavioral studies

Our model representation (Φ) of AM illusion climbed to a peak at around 3 Hz and then dropped (e.g. Fig. 3D), similar to reports for AM perception in (Sanders et al., 2013), where individuals with schizophrenia perceived lower illusion and the peak of the illusion occurred when presentation frequency of AM stimulus was at 3.12Hz. Behavioral studies (Anderson et al., 2017;Silverstein et al., 2017) suggested that individuals with schizophrenia have impaired lateral connectivity in V1, which suggests reduced inhibition. In Fig. 3D, the plot of the illusion representation Φ moved downward as the inhibitory width *σ_Inh_* decreased, just as the percept-frequency plot for schizophrenia shifting down in comparison to healthy controls (see Figure 1a in (Sanders et al., 2013)). While our results are consistent with these previous studies, we took an additional step to provide mechanistic support for the hypothesis that reduced lateral inhibition can cause impaired representation of AM illusion.

Moreover, our estimation of the illusion representation for contrast-contrast stimulus is also consistent with perceptual data. The model representation *r* for contrast-contrast illusion decreased as the inhibitory width *σ_Inh_* decreased as shown in Fig. 3A, in line with (Dakin et al., 2005), who reported that individuals with schizophrenia perceive weaker contrast-contrast illusion than healthy controls. Our findings again provide additional mechanistic support to the hypothesis that reduced lateral inhibition could cause impaired contrast-contrast illusion.

The changing patterns of perception for AM and contrast-contrast illusions have not been studied comparatively. Here we found that the changing pattern of the model representation across different values of *σ_Inh_* for AM illusion simulation was the same as for contrast-contrast illusion (see the curve shapes in Fig. 3C and Fig. 3A). These findings support and validate the use of our proposed two-layer model for probing neurobiological changes in conditions such as schizophrenia and their impact on visual percepts.

Our results are consistent with reports that individuals with schizophrenia have reduced top-down feedback (Gilbert and Sigman, 2007;Silverstein, 2016) and reveal a plausible mechanism showing that reduced top-down feedback can impair contrast-contrast and AM illusions percept. Furthermore, our results showed that the model representation for both contrast-contrast and AM illusions shared the same trend when the feedback strength *α* decreased (see the plots in Fig. 4D and Fig. 4A).

Our results are also in line with reports that individuals with schizophrenia have increased lateral excitation (Silverstein et al., 2017), suggesting a mechanism, through which increasing the excitatory width *σ_EX_* in a range of higher values can cause impaired contrast-contrast and AM illusions. In addition, our results revealed that the model representation trends were the same for both illusions when the excitatory width (*σ_Ex_*) increased within a range of higher values (see the right panels in Figs. 5A&C).

### 4.2 Top-down feedback can enlarge the difference in illusion representation

In our model, the simulated illusion representation did not change linearly with the lateral inhibition change. The simulated illusion representation increased as the width of the inhibitory subfield got larger, but the change rate of the illusion representation became smaller. For instance, Fig. 8A shows that the model illusion representation decreased slowly between the two purple “x” marks. In this range, the change of the illusion representation may not be large enough to be interpreted as a significant change of illusion representation in higher association cortices. This suggests that impaired lateral inhibition alone in schizophrenia (Anderson et al., 2017) might not be enough to explain incongruent findings, e.g., why Dakin et al. (Dakin et al., 2005) reported significant difference between individuals with schizophrenia and controls in perceiving the contrast-contrast illusion, while Barch et al. (Barch et al., 2012) did not. This supports the hypothesis that there might be considerable heterogeneity among individuals with schizophrenia (Table 2). Because lateral inhibition is a mechanism underlying low-level visual integration our findings can also explain decreased susceptibility to some low-level integration of visual illusions in schizophrenia (King et al., 2017). Based on this, it is likely that disruption of top–down perceptual organization mechanisms can be the major factor that a more consistent resistance to high-level illusions was found in individuals with schizophrenia (King et al., 2017).

**Table 2.** Possible combined effects of parameter changes on model representation of illusions that can be used to identify distinct types of SZ across a heterogeneous spectrum. *α* is the feedback strength in the two-layer model. *σ_Inh_* is the width of the inhibitory subfield. *σ_Ex_* is the width of the excitatory subfield.

| Group | $\sigma_{Inh}$ | $\alpha$ | $\sigma_{Ex}$ | Impairment | Model representation of illusion |
| --- | --- | --- | --- | --- | --- |
| Controls | 2 | 0.3 | 0.75 | None | No reduction |
| SZ category 1 | 2 | 0.2 | 0.75 | Decreased $\alpha$ | Large reduction |
| SZ category 2 | 1.75 | 0.3 | 0.75 | Decreased $\sigma_{Inh}$ | Reduction |
| SZ category 3 | 1.75 | 0.2 | 0.75 | Decreased $\sigma_{Inh}$, decreased $\alpha$ | Larger reduction |
| SZ category 4 | 1.75 | 0.3 | 1.25 | Decreased $\sigma_{Inh}$, increased $\sigma_{Ex}$ | More reduction compared to SZ category 2 |
| SZ category 5 | 1.75 | 0.3 | 0.25 | Decreased $\sigma_{Inh}$, decreased $\sigma_{Ex}$ | More reduction compared to SZ category 2 |

Indeed, changing the inhibition and feedback simultaneously produced much larger changes in illusion representation. For example, the blue dashed line between the two purple “x” marks in the sketch in the Fig. 8B shows a relatively smaller reduction in illusion, when decreasing the width of the inhibitory subfield alone, compared to the brown oblique dashed line between the red “x” mark and the purple “x” mark that shows a greater reduction in illusion when decreasing the width of the inhibitory subfield and the top-down feedback simultaneously (see also Figs. 4C&F, summarized in sketch of Fig. 8B).

Notredame et al. (Notredame et al., 2014) proposed a Bayesian framework that combines sensory evidence and prior knowledge to explain visual perception changes in schizophrenia. Dima, Roiser, et al. (Dima et al., 2009) found that during the hollow-mask illusion task the neurophysiological signals from high-order, association cortices to lower-in-the-visual-hierarchy cortices were significantly different between individuals with schizophrenia and controls and correlated the reduction of illusion perception with weakened top-down modulation. Behavioral and neurophysiological studies for AM illusion (Sanders et al., 2012;Edwards et al., 2017) also suggest there is correlation between reduction of AM illusion and weakened feedback from higher association cortices to V1 in schizophrenia. These studies inferred that the perceived illusion was influenced by the stored information of experience (prior knowledge) passed from higher association cortices to lower cortices. That is to say that the Bayesian model explaining visual perception in schizophrenia (Notredame et al., 2014) is based on the correlation relationship between prior knowledge and feedforward visual input.

In contrast, our two-layer network provides a mechanistic relationship between weakened top-down feedback and reduced representation of illusions (Fig. 8B). The top-down feedback in our two-layer network is a mechanism that does not include any stored information of experience (priors). This suggests that the decreased top-down feedback can cause the reduced AM illusion without any top-down process of prior knowledge, which is in line with a previous study that showed no significant changes in the formation of priors in individuals with schizophrenia (Kaliuzhna et al., 2019). Our simulation results show that the decreased top-down feedback from a higher area to a lower one (cortex or thalamus) can amplify the reduction of the illusion representation.

In our neural model, impaired lateral inhibition reduced the representation of illusions too, and the reduction was magnified by weakened top-down feedback (Figs. 4C&F, 8B). Therefore, we propose that depending on the level of weakening of the feedback, there would be variable changes in illusion perception in individuals with schizophrenia. This suggests that there may be a range of percept change of illusions in schizophrenia, depending on the extent of top-down feedback reduction. As mentioned, the top-down feedback is a magnifying factor in perceiving illusions: the top-down feedback regulates illusion perception more coarsely, whereas lateral inhibition impact is finer (Figs. 4C&F, 8B). Our findings suggest that the variations in the inhibitory surround or the excitatory center can replicate minor differences of illusion precepts between individuals with schizophrenia and neurotypical controls, whereas the top-down feedback can enlarge and magnify these differences in the illusion perception.

### 4.3 Implications for diagnostic and therapeutic approaches

Based on our simulation results, it might be possible to classify people with schizophrenia into several categories with variable levels of reduced illusion representation (Fig. 8): (1) patients with impaired top-down feedback; (2) patients with impaired width of inhibitory subfields (reduced GABA); (3) patients with a combination of reduced top-down feedback and width of inhibitory subfields; (4) patients with increased width of excitatory subfields (e.g., overactive dopamine); and (5) decreased width of excitatory subfields (e.g., reduced dopamine). The last two categories can be combined as patients with either increased or decreased excitatory field outside an optimal mid-range (Fig. 8C). Table 2 shows how the changes of parameters can influence the illusion representation when they deviate from scaled values *σ_Inh_* = 2, *α* = 0.3 and *σ_Ex_* = 0.75, which are for typical responses (see the group “Controls”, first row in Table 2). The influencing factors (parameters) may change simultaneously and form a spectrum that includes additional combinations of the above scenarios, which could reveal the underlying source of heterogeneity in illusion perception in individuals with schizophrenia (Fig. 8E). For example, changes in feedback *α* alone, or in combination with changes in the inhibitory subfields (SZ categories 1 and 3, Table 2), show progressively larger reduction of illusion representation compared to the other parameter sets in Table 2. The last two rows of Table 2 show that reducing the width of the inhibitory subfield while decreasing or increasing the width of the excitatory subfield can also lead to a reduction in illusion perception. Grouping individuals with schizophrenia in categories, as suggested here, when analyzing individual differences in illusion perception may help interpret behavioral and other disease heterogeneity.

A more nuanced classification of SZ can improve our understanding of mechanisms underlying treatment approaches. Experimental approaches that can directly measure the pathophysiological underpinnings of SZ are traditionally used for the identification of biomarkers (Kraguljac et al., 2021), but theoretical and computational studies like this can further help disambiguate contradictory findings and lead to development of effective, validated biomarkers (Murray and Anticevic, 2017). For example, dopamine antagonists are not effective for all patients (McCutcheon et al., 2020). Decreasing dopamine can lead to normalized behaviors, as in healthy controls, but often this approach has no effects. This can be explained in our model, in which decreasing dopamine is reflected by decreasing the width of the excitatory subfield. Figs. 5A&C and 8C-E show that decreasing the width of the excitatory subfield does not always lead to increased illusion representation: only within an optimal mid-range of the excitatory subfield (i.e., a scaled value of *σ_Ex_* in range [0.75-1.5]) we observe increased illusion representation, which is a behavior in healthy controls, whereas a smaller excitatory width (i.e., a scaled value of *σ_Ex_* in range [0.25-0.75]) does not lead to increased illusion representation.

### 4.4 Conclusion

In this work, we proposed a two-layer network model, applied it to simulate neural representation of contrast-contrast and apparent motion illusions and observed similar results between the simulations of the two illusions. The results were also consistent with behavioral findings in previous studies with human subjects (Dakin et al., 2005;Sanders et al., 2013).

By searching the parameter space of the model and also changing each parameter independently, we found that illusion representation reduced as the width of the inhibitory surround decreased, which likely reflects impaired lateral connectivity in individuals with schizophrenia. We also found that reduced illusion representation can be caused by the decrement of the top-down feedback or the change of the width in the excitatory subfield outside an optimal mid-range.

Key contributions of our model include providing evidence for a mechanism that explains how top-down feedback can enlarge the change of illusion representation and support for the hypothesis that illusion perception can be affected by the top-down feedback without the need for stored contextual information or experience (priors). Our model shows that decreasing the top-down feedback and decreasing the width of the inhibitory subfield simultaneously amplifies reduction in illusion representation compared to decreasing the width of the inhibitory subfield alone. This can explain contradictory observations in previous studies and explain heterogeneity in SZ, reflected through variability in the severity of SZ-like illusion representation, depending on whether the top-down feedback reduction is large or small.

## 6 Author Contributions

JZ: design, methodology, formal analysis and writing of first draft. BZ and AY: funding acquisition, design, methodology, and resources. JZ, BZ, and AY: writing. All authors participated in the conceptualization of this work and approved the submitted version.

## 8 Acknowledgments

## Acknowledgements

We would like to thank Sangwook Park, Natalia Matuk, Megan Qin Deng, Cici Chen, and Caroline Dugan for their useful comments during discussions of this work.

## 10 Tables

## 11 Figure Captions

## 12 Supplementary Material

None

## 12 Data Availability Statement

The source codes for the neural model and analysis are available online (https://github.com/JiatingZhu/2LMotifVisNet)

